# Synthesis and in vitro evaluation shows disquaramide compounds are a promising class of anti-leishmanial drugs

**DOI:** 10.1101/2024.08.23.605637

**Authors:** Hayden Roys, Assyl Arykbayeva, Sarah K. Friedman, Gabriel Gifford, Eva R. Palmer, Alex Rogers, Emily N. H. Tran, Lucy Fry, Alexx Weaver, Anne Bowlin, Meghan D. Jones, Michael R. Eledge, Karl W. Boehme, Gregory R. Naumiec, Tiffany Weinkopff

## Abstract

An increasing number of treatment failures with current pharmaceutics, as well as a lack of a vaccine, demonstrates the need to develop new treatment options for leishmaniasis. Herein, we describe the synthesis and in vitro analysis of 24 disquaramide compounds targeting the *Leishmania major* parasite. Of the compounds that were evaluated, six of them (**13**, **19**, **20**, **22**, **24**, and **26**) were capable of significantly decreasing the number of parasites by up to 42% compared to the control by day four. This demonstrates that disquaramides either impair parasite replication or have leishmancidal effects. Additionally, none of the disquaramide compounds tested displayed host cell cytotoxicity. These experiments provide evidence that disquaramides have the potential to be effective anti-leishmanial therapeutics.

## INTRODUCTION

Leishmaniasis is a neglected tropical disease that affects 12 million people infected worldwide with an estimated 1.5 million new cases annually ^1^. Various species from the genus *Leishmania* infect their hosts through the bite of phlebotomine sandflies. *Leishmania* is a major global health problem with the largest impact in the eastern Mediterranean Basin, Asia, and the Americas ^2^. Leishmaniasis is a complex disease characterized by the interaction between the obligate intracellular parasite and the host’s immune response; the clinical disease can present as a wide variety of manifestations including the following categories: cutaneous leishmaniasis (CL), mucocutaneous leishmaniasis (MCL), diffuse or disseminated leishmaniasis (DCL), and visceral leishmaniasis (VL). CL, MCL, and DCL are characterized by localized lesions in the skin and/or mucosal surfaces whereas VL is a potentially life-threatening condition in which parasites migrate to internal organs including the liver, spleen, and bone marrow ^3^. Without treatment, VL is fatal. CL can result in non-healing lesions, permanent disfigurement, and disability which have long-lasting psycho-social effects ^4^. Despite being a major public health concern, there is currently no human vaccine available to protect against *Leishmania* infection.

Clinical management of leishmaniasis is largely dependent on chemotherapy using pentavalent antimonial compounds ^5^. Unfortunately, antimonial compounds have a small therapeutic window, and long treatment periods, and have demonstrated serious toxicity to the heart, liver, and pancreas of patients leading to serious adverse effects including vomiting, myalgia, pain, lethargy, and in rare cases, fatal cardiac arrhythmia ^6^. Moreover, in high-risk groups, including pregnant women, the risks of antimonial treatment can outweigh the benefits ^4^. Due to widespread use, antimonial resistance has also become a major issue in *Leishmania*-endemic regions ^7^. In light of antimonial resistance, miltefosine has been licensed as a second-line treatment for many forms of leishmaniasis. Miltefosine is promising because it is the first oral treatment for leishmaniasis; all others are delivered by the intraveneous route. However, there are important concerns regarding patients of reproductive age due to the embryo-fetal toxicity associated with miltefsine treatment. Due to embryo-fetal toxicity, patients must receive effective preventive contraception throughout anti-parasitic treatment and for an additional five months following treatment ^4, 8^. Additionally, patients frequently report gastrointestinal issues and other negative side effects while taking miltefsine ^4^. In recent years, there have been a few reports of miltefosine-resistant *Leishmania* detected in the clinic ^7^. Amphotericin B (AmB) is a polyene natural product originally formulated as an anti-fungal treatment that has been used as a treatment for leishmaniasis ^9^. To reduce toxicity, AmB was converted into a liposomal formulation to treat VL. Unfortunately, there have also been reported instances of AmB resistance and treatment failure in leishmaniasis patients ^9–11^. Given the ongoing health crisis of leishmaniasis, and the alarming increase in resistance, there is a need for new anti-leishmanial compounds.

A structural class of compounds known as disquaramides has shown promise against *Leishmania* parasites ^12^. Disquaramides exhibit known anti-parasitic properties, with low human toxicity, and are relatively simple and inexpensive to synthesize ^12, 13^. They are known to damage the cellular structures of the *Leishmania* promastigotes, characterized by rupture of the mitochondrial membrane and damage to the outer membrane of the parasite which negatively affects other organelles ^12^. This results in parasite immobility and/or death ^10^. Currently, only a handful of compounds in this structural class have been examined for their leishmanicidal properties, and of them, one is nearly as effective as glucantime in regard to leishmanicidal activities ^12^. Of the 24 disquaramides successfully synthesized and reported below, several candidates of this structural class show potential for a potent, inexpensive, and safe drug for leishmaniasis. These results find disquaramide compounds are effective anti-leishmanial agents against both extracellular parasites as well as parasites residing in infected macrophages.

## MATERIALS AND METHODS

### Mice

Female C57BL/6NCr mice were purchased from the National Cancer Institute and housed in the Division of Laboratory Animal Medicine at the University of Arkansas for Medical Sciences (UAMS) under pathogen-free conditions. Mice between 6 and 8 weeks of age were used to prepare macrophages from the bone marrow. All procedures were performed in accordance with the guidelines of the UAMS Institutional Animal Care and Use Committee (IACUC).

### Parasites and infections

Fluorescently-labeled DsRed *Leishmania major* (*L. major*) (WHO/MHOM/IL/80/Friedlin) parasites were maintained in vitro in Schneider’s Drosophila medium (Gibco) supplemented with 20% heat-inactivated FBS (Invitrogen), 2 mM L-glutamine (Sigma), 100 U/mL penicillin, and 100 mg/mL streptomycin (Sigma). For the direct culture of parasites with the compounds (in the absence of macrophages), promastigotes were isolated by centrifugation at day 4. For the culture of infected macrophages with compounds, the metacyclic promastigote stage of parasites was isolated from cultures at day 4 by Ficoll (Sigma) gradient separation ^14^.

### Synthesis of disquaramide compounds

Unless stated otherwise, all reactions were carried out in oven-dried glassware. NMR spectra were obtained using a JEOL ECX-300 spectrometer (Peabody, MA) at 300 MHz (^1^H NMR) and 75 MHz (^13^C NMR). Chemical shifts are in ppm relative to the CDCl_3_ or DMSO-d_6_ resonance. Spin-spin coupling constants (*J*) are given in Hz. Mass spectra were obtained using a Waters MALDI micro MX TOF spectrometer (Milford, MA) matrix-free. Purity was assessed using a Shimadzu LC-2050 analytical HPLC system (Columbia, MD). Analytical TLC was performed on Sorbtech (Norcross, GA) polyester-backed TLC plates (TLC silica gel 60 UV254) and compounds were detected with a UV lamp. Flash chromatography was performed using a Teledyne ISCO CombiFlash NG 100 (Lincoln, NE) with Redisep Silver normal phase silica gel (40-60 μm, 230-400 mesh). All reagents and solvents were used as is, without further purification. All reagents were purchased from Fisher Scientific, except for 3,4-diethoxy-3-cyclobutene-1,2-dione (CombiBlocks).

### Synthesis of monosquaramides

#### 3-(Butylamino)-4-ethoxycyclobut-3-ene-1,2-dione (2)

To a stirred solution of 3,4-diethoxycyclobut-3-ene-1,2-dione (0.85 g, 5.0 mmol) in 200 proof ethanol (5 mL) was added a solution of *n*-butylamine (0.49 mL, 5.0 mmol) in 200 proof ethanol (10 mL) dropwise over 10 min. Once the addition was complete, the reaction was allowed to stir for 2 hours at room temperature (RT). The reaction mixture was then concentrated under reduced pressure and the resulting crude solid was resuspended in hexanes. The solid was collected by vacuum filtration without any further purification to afford **2**, a white solid, in 86.6% yield (0.853 g). All spectroscopic data agrees with previously reported literature values ^13^.

#### 3-((3-(dimethylamino)propyl)(methyl)amino)-4-ethoxycyclobut-3-ene-1,2-dione (3)

To a stirred solution of 3,4-diethoxycyclobut-3-ene-1,2-dione (0.85 g, 5.0 mmol) in 200 proof ethanol (5 mL) was added a solution of *N*^1^*,N*^1^*,N*^3^*-trimethylpropane-1,3-diamine* (0.73 mL, 5.0 mmol) in 200 proof ethanol (10 mL) dropwise over 10 min. Once the addition was complete, the reaction was allowed to stir for 2 h at RT. The reaction mixture was then concentrated under reduced pressure. The resulting crude oil was purified by flash chromatography (SiO_2_, 100% EtOAc → 95% EtOAc/ 5%MeOH) to afford **3**, a pale yellow oil, in 83.7% yield (1.008 g). All spectroscopic data agrees with previously reported literature values ^13^.

### Synthesis of disquaramides

#### General procedure A

Compounds **4−14 and 17** were synthesized using the following general procedure. To a stirred solution of **2** (0.183 g, 1.0 mmol) in 200 proof ethanol (5 mL) was added a solution of amine (2.0 mmol) in 200 proof ethanol (10 mL) dropwise over 10 min. Once the addition was complete, the reaction was allowed to stir for 2−16 h at RT. The reaction mixture was then concentrated under reduced pressure and the resulting crude product was either resuspended in hexane and collected by filtration without further purification or purified by flash chromatography (SiO_2_, 100% EtOAc → 95% EtOAc/ 5%MeOH).

#### General Procedure B

Compounds **15**, **16**, **and 18−27** were synthesized using the following general procedure. To a stirred solution of **3** (0.240 g, 1.0 mmol) in 200 proof ethanol (5 mL) was added a solution of amine (2.0 mmol) in 200 proof ethanol (10 mL) dropwise over 10 min. Once the addition was complete, the reaction was allowed to stir for 2−16 h at RT. The reaction mixture was then concentrated under reduced pressure and the resulting crude product was either resuspended in hexane and collected by filtration without further purification or purified by flash chromatography (SiO_2_, 100% EtOAc → 95% EtOAc/ 5%MeOH).

#### 3-(butylamino)-4-(propylamino)cyclobut-3-ene-1,2-dione (4)

**Using General Procedure A**, compound **4** was synthesized using *n*-propylamine (0.16 mL, 2.0 mmol). The reaction mixture was stirred at RT for 2 h and the crude solid was resuspended in hexanes and isolated by vacuum filtration without further purification to afford **4**, a white solid, in 80.4% yield (0.169 g). ^1^H NMR (300 MHz, DMSO-d_6_):δ 7.31 (br s, 2H),3.49−3.45 (m, 4H), 1.53−1.45 (m, 4H), 1.30 (sextet, *J* = 7.56 Hz, 2H), 0.89−0.86 (m, 6H). ^13^C NMR (75 MHz, DMSO-d_6_): δ 182.3 (2C), 167.8 (2C), 49.9, 42.9, 32.9, 24.0, 19.0, 13.5, 10.8. HRMS (MALDI) m/z calculated for C_11_H_18_N_2_O_2_, 210.1368 g/mol; found, 210.1660 g/mol. Purity >99%.

#### 3-(butylamino)-4-(isopropylamino)cyclobut-3-ene-1,2-dione (5)

**Using General Procedure A**, compound **5** was synthesized using isopropylamine (0.18 mL, 2.0 mmol). The reaction mixture was stirred at RT for 2 h and the crude solid was resuspended in hexanes and isolated by vacuum filtration without further purification to afford **5**, a white solid, in 77.6% yield (0.163 g). ^1^H NMR (300 MHz, DMSO-d_6_): δ 7.30 (br s, 2H), 4.08 (septet, *J* = 6.18 Hz, 1H), 3.49 (m, 2H), 1.48 (quintet, *J* = 7.23 Hz, 2H), 1.30 (sextet, *J* = 7.20 Hz, 2H), 1.17 (d, *J* = 6.54 Hz, 6H), 0.88 (t, *J* = 7.23 Hz, 3H). ^13^C NMR (75 MHz, DMSO-d_6_): δ 182.2, 182.0, 167.8, 167.0, 45.5, 42.9, 32.8, 23.8 (2C), 19.0, 13.5. HRMS (MALDI) m/z calculated for C_11_H_18_N_2_O_2_, 210.1368 g/mol; found, 210.1660 g/mol. Purity >99%.

#### 3,4-bis(butylamino)cyclobut-3-ene-1,2-dione (6)

Using General Procedure A, compound **6** was synthesized using *n*-butylamine (0.20 mL, 2.0 mmol). The reaction mixture was stirred at RT for 12 h and the crude solid was resuspended in hexanes and isolated by vacuum filtration without further purification to afford **6**, a white solid, in 86.4% yield (0.194 g). All spectroscopic data agrees with previously reported literature values ^15^.

#### 3-(butylamino)-4-(isobutylamino)cyclobut-3-ene-1,2-dione (7)

Using General Procedure A, compound **7** was synthesized using isobutylamine (0.20 mL, 2.0 mmol). The reaction mixture was stirred at RT for 2 h and the crude solid was resuspended in hexanes and isolated by vacuum filtration without further purification to afford **7**, a white solid, in 59.0% yield (0.132 g). ^1^H NMR (300 MHz, DMSO-d_6_): δ 7.29 (br s, 2H), 3.89−3.40 (m, 2H), 3.50−3.48 (m, 2H), 1.52−1.44 (m, 3H), 1.30 (sextet, *J* = 7.53 Hz, 2H), 1.63−1.41 (m, 3H), 0.90−0.82 (m, 6H). ^13^C NMR (75 MHz, DMSO-d_6_): δ 182.1, 182.0, 167.7, 167.3, 50.9, 42.9, 32.8, 30.1, 21.6, 19.0, 13.5, 10.1 (2C). HRMS (MALDI) m/z calculated for C_12_H_20_N_2_O_2_, 224.1525 g/mol; found, 224.1817 g/mol. Purity >99%.

#### 3-(butylamino)-4-(pentylamino)cyclobut-3-ene-1,2-dione (8)

Using General Procedure A, compound **8** was synthesized using *n*-pentylamine (0.24 mL, 2.0 mmol). The reaction mixture was stirred at RT for 6 h and the crude solid was resuspended in hexanes and isolated by vacuum filtration without further purification to afford **8**, a white solid, in 86.4% yield (0.158 g). ^1^H NMR (300 MHz, DMSO-d_6_): δ 7.29 (br s, 2H), 3.47 (br s, 4H), 1.52−1.42 (m, 4H), 1.34−1.25 (m, 6H), 0.89−0.84 (m, 6H). ^13^C NMR (75 MHz, DMSO-d_6_): δ 182.2 (2C), 167.7 (2C), 43.2, 42.9, 32.9, 30.5, 28.0, 21.8, 19.0, 13.9, 13.5. HRMS (MALDI) m/z calculated for C_13_H_22_N_2_O_2_, 238.1681 g/mol; found, 238.1973 g/mol. Purity >99%.

#### 3-(butylamino)-4-(tert-pentylamino)cyclobut-3-ene-1,2-dione (9)

Using General Procedure A, compound **9** was synthesized using 2-methylbutan-2-amine (0.24 mL, 2.0 mmol). The reaction mixture was stirred at RT for 12 h and the crude solid was resuspended in hexanes and isolated by vacuum filtration without further purification to afford **9**, a white solid, in 57.1% yield (0.137 g). ^1^H NMR (300 MHz, DMSO-d_6_): δ 7.48 (br s, 1H), 7.38 (br s, 1H), 3.52 (q, *J* = 6.18 Hz, 2H), 1.67 (sextet, *J* = 7.56 Hz, 2H), 1.51 (pentet, *J* = 7.23 Hz, 2H), 1.34−1.29 (m, 8H), 0.88 (t, *J* = 7.20 Hz, 3H), 0.83 (t, *J* = 7.20 Hz, 3H). ^13^C NMR (75 MHz, DMSO-d_6_): δ 182.2, 180.8, 168.6, 167.7, 54.8, 43.0, 34.4, 32.7, 27.6 (2C), 19.0, 13.5, 8.4. HRMS (MALDI) m/z calculated for C_13_H_22_N_2_O_2_, 238.1681 g/mol; found, 238.1973 g/mol. Purity >99%.

#### 3-(benzylamino)-4-(butylamino)cyclobut-3-ene-1,2-dione (10)

Using General Procedure A, compound **10** was synthesized using benzylamine (0.22 mL, 2.0 mmol). The reaction mixture was stirred at RT for 2 h and the crude solid was resuspended in hexanes and isolated by vacuum filtration without further purification to afford **10**, a white solid, in 77.9% yield (0.201 g). ^1^H NMR (300 MHz, DMSO-d_6_): δ 7.73 (br s, 1 H), 7.36−7.32 (m, 6H), 4.69 (d, *J* = 5.82 Hz, 2H), 3.49−3.47 (m, 2H), 1.47 (pentet, *J* = 7.53 Hz, 2H), 1.28 (sextet, *J* = 7.53 Hz, 2H), 0.86 (t, *J* = 7.23 Hz, 3H). ^13^C NMR (75 MHz, DMSO-d_6_): δ 182.6, 182.3, 168.0, 167.3, 139.0, 128.6 (2C), 127.5 (2C), 127.4, 46.8, 43.0, 32.8, 19.0, 13.5. HRMS (MALDI) m/z calculated for C_15_H_18_N_2_O_2_, 258.1368 g/mol; found, 258.1660 g/mol. Purity >99%.

#### 3-(butylamino)-4-(diethylamino)cyclobut-3-ene-1,2-dione (11)

Using General Procedure A, compound **11** was synthesized using diethylamine (0.22 mL, 2.0 mmol). The reaction mixture was stirred at RT for 4 h and the crude solid was resuspended in hexanes and isolated by vacuum filtration without further purification to afford **11**, a white solid, in 83.4% yield (0.187 g). ^1^H NMR (300 MHz, DMSO-d_6_): δ 7.57 (m, 1H), 3.56−3.51 (m, 6H), 1.50 (pentet, *J* = 7.23 Hz, 2H), 1.28 (sextet, *J* = 7.53 Hz, 2H), 1.12 (t, *J* = 7.20 Hz, 6H), 0.87 (t, *J* = 7.20 Hz, 3H). ^13^C NMR (75 MHz, DMSO-d_6_): δ 182.3, 181.7, 167.0, 166.7, 43.4 (2C), 43.0, 33.2, 19.0, 15.1 (2C), 13.6. HRMS (MALDI) m/z calculated for C_12_H_20_N_2_O_2_, 224.1525 g/mol; found, 224.1817 g/mol. Purity >99%.

#### 3-(butylamino)-4-((2-(dimethylamino)ethyl)amino)cyclobut-3-ene-1,2-dione (12)

Using General Procedure A, compound **12** was synthesized using *N*^1^,*N*^1^-dimethylethane-1,2-diamine (0.22 mL, 2.0 mmol). The reaction mixture was stirred at RT for 2 h and the crude solid was resuspended in hexanes and isolated by vacuum filtration without further purification to afford **12**, a white solid, in 87.9% yield (0.210 g). All spectroscopic data agrees with previously reported literature values ^13^.

#### 3-(butylamino)-4-((2-(diethylamino)ethyl)amino)cyclobut-3-ene-1,2-dione (13)

Using General Procedure A, compound **13** was synthesized using *N*^1^,*N*^1^-diethylethane-1,2-diamine (0.35 mL, 2.0 mmol). The reaction mixture was stirred at RT for 2 h and the crude solid was resuspended in hexanes and isolated by vacuum filtration without further purification to afford **13**, a white solid, in 70.0% yield (0.189 g). ^1^H NMR (300 MHz, DMSO-d_6_): δ 7.43 (br s, 1H), 7.21 (br s, 1H), 3.50−3.46 (m, 4H), 2.47−2.42 (m, 6H), 1.44 (pentet, *J* = 6.84 Hz, 2H), 1.31−1.20 (sextet, *J* = 7.20 Hz, 2H), 0.92−0.82 (m, 9H). ^13^C NMR (75 MHz, DMSO-d_6_): δ 182.4 (2C), 167.8 (2C), 53.1, 46.5 (2C), 42.8, 41.6, 32.8, 19.0, 13.5, 11.6 (2C). HRMS (MALDI) m/z calculated for C_14_H_25_N_3_O_2_, 267.1947 g/mol; found, 267.2239 g/mol. Purity >99%.

#### 3-(butylamino)-4-((3-(dimethylamino)propyl)amino)cyclobut-3-ene-1,2-dione (14)

Using General Procedure A, compound **14** was synthesized using *N*^1^,*N*^1^-dimethylpropane-1,3-diamine (0.25 mL, 2.0 mmol). The reaction mixture was stirred at RT for 2 h and the crude solid was resuspended in hexanes and isolated by vacuum filtration without further purification to afford **14**, a white solid, in 68.9% yield (0.174 g). All spectroscopic data agrees with previously reported literature values ^16^.

#### 3-((3-(dimethylamino)propyl)(methyl)amino)-4-(propylamino)cyclobut-3-ene-1,2-dione (15)

Using General Procedure B, compound **15** was synthesized using *n-*propylamine (0.16 mL, 2.0 mmol). The reaction mixture was stirred at RT for 2 h and the crude solid was resuspended in hexanes and isolated by vacuum filtration without further purification to afford **15**, a white solid, in 83.1% yield (0.123 g). ^1^H NMR (300 MHz, DMSO-d_6_): δ 7.97 (br s, 1H), 3.48 (t, *J* = 6.90 Hz, 2H), 3.39 (t, *J* = 6.60 Hz, 2H), 3.13 (s, 3H), 2.18 (t, *J* = 7.23 Hz, 2H), 2.09 (s, 6H), 1.64 (quintet, *J* = 7.20 Hz, 2H), 1.49 (sextet, *J* = 7.20 Hz, 2H), 0.83 (t, *J* = 7.23 Hz). ^13^C NMR (75 MHz, DMSO-d_6_): δ 182.3, 181.9, 167.8, 167.4, 55.1, 48.9, 45.1, 44.7 (2C), 36.1, 24.7, 24.3, 10.8. HRMS (MALDI) m/z calculated for C_13_H_23_N_3_O_2_, 253.1790 g/mol; found, 253.2082 g/mol. Purity >99%.

#### 3-((3-(dimethylamino)propyl)(methyl)amino)-4-(isopropylamino)cyclobut-3-ene-1,2-dione (16)

Using General Procedure B, compound **16** was synthesized using isopropylamine (0.18 mL, 2.0 mmol). The reaction mixture was stirred at RT for 6 h and the crude solid was resuspended in hexanes and isolated by vacuum filtration without further purification to afford **16**, a white solid, in 99.6% yield (0.253 g). ^1^H NMR (300 MHz, DMSO-d_6_): δ 7.70 (br s, 1H), 4.34 (septet, *J* = 6.90 Hz, 1H), 3.45 (br s, 2H), 3.17 (s, 3H), 2.21 (t, *J* = 6.54 Hz, 2H), 2.11 (s, 6H), 1.67 (quintet, *J* = 6.54 Hz, 2H), 1.18 (d, *J* = 6.54 Hz). ^13^C NMR (75 MHz, DMSO-d_6_): δ182.3, 181.7, 168.0, 166.6, 55.2, 48.7, 45.7, 44.9 (2C), 35.9, 24.6, 23.6 (2C). HRMS (MALDI) m/z calculated for C_13_H_23_N_3_O_2_, 253.1790 g/mol; found, 253.2082 g/mol. Purity >99%.

#### 3-(butylamino)-4-((3-(dimethylamino)propyl)(methyl)amino)cyclobut-3-ene-1,2-dione (17)

Using General Procedure A, compound **17** was synthesized using *N*^1^,*N*^1^,*N*^3^-trimethylpropane-1,3-diamine (0.29 mL, 2.0 mmol). The reaction mixture was stirred at RT for 12 h and the crude solid was resuspended in hexanes and isolated by vacuum filtration without further purification to afford **17**, a white solid, in 78.7% yield (0.210 g). All spectroscopic data agrees with previously reported literature values ^13^.

#### 3-((3-(dimethylamino)propyl)(methyl)amino)-4-(isobutylamino)cyclobut-3-ene-1,2-dione (18)

Using General Procedure B, compound **18** was synthesized using isobutylamine (0.20 mL, 2.0 mmol). The reaction mixture was stirred at RT for 2 h and the crude solid was resuspended in hexanes and isolated by vacuum filtration without further purification to afford **18**, a white solid, in 98.9% yield (0.263 g). ^1^H NMR (300 MHz, DMSO-d_6_): δ 7.99 (br s, 1H), 3.37 (t, *J* = 6.18 Hz, 2H), 3.18 (s, 3H), 2.21 (t, *J* = 6.54 Hz, 2H), 2.11 (s, 6H), 1.81−1.50 (m, 3H), 0.86 (d, *J* = 6.87 Hz, 6H). ^13^C NMR (75 MHz, DMSO-d_6_): δ 182.3, 181.9, 167.7, 167.5, 55.0, 50.5, 48.8, 44.4 (2C), 29.7, 24.4, 19.4 (2C). HRMS (MALDI) m/z calculated for C_14_H_25_N_3_O_2_, 267.1947; found, 267.2239 g/mol. Purity >99%.

#### 3-(tert-butylamino)-4-((3-(dimethylamino)propyl)(methyl)amino)cyclobut-3-ene-1,2-dione (19)

Using General Procedure B, compound **19** was synthesized using *tert*-butylamine (0.21 mL, 2.0 mmol). The reaction mixture was stirred at RT for 12 h and the crude solid was resuspended in hexanes and isolated by vacuum filtration without further purification to afford **19**, a white solid, in 48.0% yield (0.128 g). ^1^H NMR (300 MHz, DMSO-d_6_): δ 8.12 (br s, 1H), 3.49−3.32 (m, 2H), 3.21 (s, 3H), 2.26 (t, *J* = 6.18 Hz, 2H), 2.14 (s, 6H), 1.70 (quintet, *J* = 6.54 Hz, 2H), 1.38 (s, 9H). ^13^C NMR (75 MHz, DMSO-d_6_): δ 183.3, 181.0, 169.4, 167.9, 54.2, 52.1, 44.5 (2C), 35.4, 30.2 (3C). HRMS (MALDI) m/z calculated for C_14_H_25_N_3_O_2_, 267.1947 g/mol; found, 267.2239 g/mol. Purity >99%.

#### 3-((3-(dimethylamino)propyl)(methyl)amino)-4-(pentylamino)cyclobut-3-ene-1,2-dione (20)

Using General Procedure B, compound **20** was synthesized using *n*-pentylamine (0.24 mL, 2.0 mmol). The reaction mixture was stirred at RT for 2 h and the crude solid was resuspended in hexanes and isolated by vacuum filtration without further purification to afford **20**, a white solid, in 84.0% yield (0.074 g). ^1^H NMR (300 MHz, DMSO-d_6_): δ 7.93 (br s, 1H), 3.57−3.46 (m, 4H), 3.17 (s, 3H), 2.20 (t, *J* = 6.84 Hz, 2H), 2.11 (s, 6H), 1.67 (quintet, *J* = 6.84 Hz, 2H), 1.52 (quintet, *J* = 6.87 Hz, 2H), 1.28−1.25 (m, 4H), 0.86 (t, *J* = 6.87 Hz, 3H). ^13^C NMR (75 MHz, DMSO-d_6_): δ 182.2, 181.9, 167.8, 167.3, 55.1, 48.8, 44.5 (2C), 43.4, 36.1, 30.7, 28.1, 24.5, 21.8, 13.9. HRMS (MALDI) m/z calculated for C_15_H_27_N_3_O_2_, 281.2103 g/mol; found, 281.2395 g/mol. Purity >99%.

#### 3-((3-(dimethylamino)propyl)(methyl)amino)-4-(tert-pentylamino)cyclobut-3-ene-1,2-dione (21)

Using General Procedure B, compound **21** was synthesized using 2-methylbutan-2-amine (0.24 mL, 2.0 mmol). The reaction mixture was stirred at RT for 12 h and the crude solid was resuspended in hexanes and isolated by vacuum filtration without further purification to afford **21**, a white solid, in 91.9% yield (0.251 g). ^1^H NMR (300 MHz, DMSO-d_6_): δ 7.90 (br s, 1H), 3.40 (m, 2H), 3.22 (s, 3H), 2.24 (t, *J* = 4.80 Hz), 2.11 (s, 6H), 1.72−1.68 (m, 4H), 1.31 (s, 6H), 0.83 (t, *J* = 6.97 Hz, 3H). ^13^C NMR (75 MHz, DMSO-d_6_): δ 183.3, 180.9, 169.6, 167.9, 54.9, 54.3, 47.7, 44.7 (2C), 35.3, 34.3, 27.6 (2C), 22.7, 8.6. HRMS (MALDI) m/z calculated for C_15_H_27_N_3_O_2_, 281.2103 g/mol; found, 281.2395 g/mol. Purity >99%.

#### 3-(benzylamino)-4-((3-(dimethylamino)propyl)(methyl)amino)cyclobut-3-ene-1,2-dione (22)

Using General Procedure B, compound **22** was synthesized using benzylamine (0.22 mL, 2.0 mmol). The reaction mixture was stirred at RT for 2 h and the crude solid was resuspended in hexanes and isolated by vacuum filtration without further purification to afford **22**, a white solid, in 63.1% yield (0.190 g). ^1^H NMR (300 MHz, DMSO-d_6_): δ 8.41 (br s, 1H), 7.36−7.26 (m, 5H), 4.76 (d, *J* = 5.82 Hz, 2H), 3.42 (m, 2H), 3.18 (s, 3H), 2.18 (t, *J* = 6.54 Hz, 2H), 2.02 (s, 6H), 1.66 (quintet, *J* = 6.54 Hz, 2H). ^13^C NMR (75 MHz, DMSO-d_6_): δ 183.0, 182.5, 168.7, 167.5, 139.9, 129.0 (2C), 128.0 (2C), 127.8, 49.5, 47.3, 45.3. HRMS (MALDI) m/z calculated for C_17_H_23_N_3_O_2_, 301.1790 g/mol; found, 301.2082 g/mol. Purity >99%.

#### 3-(diethylamino)-4-((3-(dimethylamino)propyl)(methyl)amino)cyclobut-3-ene-1,2-dione (23)

Using General Procedure B, compound **23** was synthesized using diethylamine (0.22 mL, 2.0 mmol). The reaction mixture was stirred at RT for 12 h and the crude solid was resuspended in hexanes and isolated by vacuum filtration without further purification to afford **23**, a white solid, in 54.6% yield (0.144 g). ^1^H NMR (300 MHz, DMSO-d_6_): δ 3.60 (t, *J* = 6.18 Hz, 2H), 3.52 (q, *J* = 6.87 Hz, 4H), 3.12 (s, 3H), 2.23 (t, *J* = 6.87 Hz, 2H), 2.13 (s, 6H), 1.72 (quintet, *J* = 7.20 Hz. 2H), 1.15 (t, *J* = 7.23 Hz, 6H). ^13^C NMR (75 MHz, DMSO-d_6_): δ 183.7, 183.5, 169.7, 168.7, 56.4, 50.8, 45.5 (2C), 44.8 (2C), 39.3, 25.8, 14.4 (2C). HRMS (MALDI) m/z calculated for C_14_H_25_N_3_O_2_, 267.1947 g/mol; found, 267.2239 g/mol. Purity >99%.

#### 3-(dibutylamino)-4-((3-(dimethylamino)propyl)(methyl)amino)cyclobut-3-ene-1,2-dione (24)

Using General Procedure B, compound **24** was synthesized using dibutylamine (0.34 mL, 2.0 mmol). The reaction mixture was stirred at RT for 12 h and the crude oil was purified by flash chromatography (100% EtOAc → 95% EtOAc/5% MeOH to afford **24**, an amber oil, in 79.4% yield (0.201 g). ^1^H NMR (300 MHz, DMSO-d_6_): δ 3.60 (t, *J* = 7.20 Hz, 2H), 3.48 (t, *J* = 7.20 Hz, 4H), 3.11 (s, 3H), 2.19 (t, *J* = 6.90 Hz, 2H), 2.11 (s, 6H), 1.70 (quintet, *J* = 6.84 Hz, 2H), 1.54 (quintet, *J* = 6.54 Hz, 4H), 1.25 (sextet, *J* = 7.20 Hz, 4H), 0.88 (t, *J* = 7.23 Hz, 6H). ^13^C NMR (75 MHz, DMSO-d_6_): δ 183.4, 183.1, 169.3, 168.9, 55.9, 50.2, 49.5 (2C), 45.0 (2C), 38.9, 30.2 (2C), 25.3, 19.2 (2C), 13.6 (2C). HRMS (MALDI) m/z calculated for C_18_H_33_N_3_O_2_, 323.2573 g/mol; found, 323.2865 g/mol. Purity >99%.

#### 3-((2-(dimethylamino)ethyl)amino)-4-((3-(dimethylamino)propyl)(methyl)amino)cyclobut-3-ene-1,2-dione (25)

Using General Procedure B, compound **25** was synthesized using *N*^1^,*N*^1^-dimethylethane-1,2-diamine (0.22 mL, 2.0 mmol). The reaction mixture was stirred at RT for 12 h and the crude solid was resuspended in hexanes and isolated by vacuum filtration without further purification to afford **25**, a yellow solid, in 97.2% yield (0.274 g). ^1^H NMR (300 MHz, DMSO-d_6_): δ 7.89 (br s, 1H), 3.62 (q, *J* = 6.18 Hz, 2H), 3.41 (m, 2H), 3.17 (s, 3H), 2.38 (t, *J* = 6.51 Hz, 2H), 2.20 (t, *J* = 6.87 Hz, 2H), 2.15 (s, 6H), 2.11 (s, 6H), 1.67 (quintet, *J* = 6.54 Hz, 2H). ^13^C NMR (75 MHz, DMSO-d_6_): δ 182.3, 181.9, 167.9, 167.3, 59.8 (2C), 55.0, 48.9, 45.3 (2C), 44.9, 41.5, 36.1, 24.7. HRMS (MALDI) m/z calculated for C_14_H_26_N_4_O_2_, 282.2056 g/mol; found, 282.2348 g/mol. Purity >99%.

#### 3-((2-(diethylamino)ethyl)amino)-4-((3-(dimethylamino)propyl)(methyl)amino)cyclobut-3-ene-1,2-dione (26)

Using General Procedure B, compound **26** was synthesized using *N*^1^,*N*^1^-diethylethane-1,2-diamine (0.35 mL, 2.0 mmol). The reaction mixture was stirred at RT for 12 h and the crude oil was purified by flash chromatography (100% EtOAc → 95% EtOAc/5% MeOH) to afford **26**, a yellow oil, in 65.4% yield (0.200 g). ^1^H NMR (300 MHz, DMSO-d_6_): δ 7.81 (br s, 1H), 3.51−3.36 (m, 4H), 3.10 (s, 3H), 2.46−2.38 (m, 6H), 2.18−2.06 (m, 8H), 1.61 (quintet, *J* = 6.87 Hz, 2H), 0.86 (t, *J* = 6.87 Hz, 6H). ^13^C NMR (75 MHz, DMSO-d_6_): δ 182.3, 181.9, 167.8, 167.4, 55.0, 53.3, 48.8, 46.8 (2C), 44.8 (2C), 41.8, 36.0, 24.8, 11.9 (2C). HRMS (MALDI) m/z calculated for C_16_H_30_N_4_O_2_, 310.2369 g/mol; found, 310.2661 g/mol. Purity >99%.

#### 3-((3-(dimethylamino)propyl)(methyl)amino)-4-((3- (dimethylamino)propyl)amino)cyclobut-3-ene-1,2-dione (27)

Using General Procedure B, compound **27** was synthesized using *N*^1^,*N*^1^-dimethylpropane-1,3-diamine (0.25 mL, 2.0 mmol). The reaction mixture was stirred at RT for 12 h and the crude solid was resuspended in hexanes and isolated by vacuum filtration without further purification to afford **27**, a white solid, in 86.1% yield (0.254 g). All spectroscopic data agrees with previously reported literature values ^17^.

### Antileishmanial activity in vitro assay

To test the direct effects of disquaramide compounds on parasites, 5×10^6^ *L. major* DsRed promastigote parasites were cultured in T25 flasks in 10 mL of complete Schneider’s media (Gibco) with or without disquaramide compounds for 4 days at concentrations of 1 μM, 5 μM, or 10 μM in triplicate. On each day, 10 μL of parasite-containing media from each flask was added to 90 μL of 2% PFA (EMS) to fix and dilute the parasites at a 1:10 ratio in a 96-well flat bottom plate. The plate was imaged using the 10X wide field objective on a BioTek Cytation C10 Confocal Imaging Reader in the TxRed channel. Biotek Gen5 software was used to quantify parasites.

### Generation of bone marrow-derived macrophages

Bone marrow-derived macrophages (BMDM) were cultured in Dulbecco’s modified Eagle medium (DMEM, Gibco) supplemented with 10% FBS (Invitrogen), 2.5% HEPES (Gibco), 1% antibiotics (pen/strep, Sigma), and 25% supernatant from L929 cells as a source of M-CSF for 7 days. For experiments, adherent macrophages were removed after 7 days, counted, and transferred to plates in DMEM to allow for adherence overnight at 37 °C before in vitro infection or exposure to drug compounds.

### Antileishmanial activity in vitro macrophage assay

BMDM (1×10^6^ cells per well) were infected with DsRed fluorescent *L. major* parasites (MOI 5:1) for 2 hours at 37 °C in a 24-well plate in DMEM to allow parasite internalization. After 2 hours of infection, BMDM were washed 3 times to remove extracellular parasites. Next, BMDM were cultured for 72 h in the presence or absence of different drug treatments. Specifically, BMDM were treated with various disquaramide compounds at 3 different concentrations (1 μM, 5 μM, or 10 μM) during the infection. BMDM were cultured with disquaramide compounds for the duration of the experiment after extracellular parasites were washed away. For a positive control to induce parasite death, BMDM were cultured with 1 μM, 5 μM, or 10 μM miltefosine (Sigma). After 72 h BMDM were fixed with ice-cold methanol for 3 minutes and washed twice with PBS (Gibco); nuclei were then stained with DAPI (Invitrogen). Excess DAPI was removed and washed 3 times before the cells were submerged in 500 μL PBS for imaging.

Imaging was acquired on a Cytation 10 confocal imager and analyzed with BioTek Gen5 software that allows for parasite and macrophage cell counting. To calculate the number of parasites per macrophage, the number of parasites was divided by the number of macrophages. Experiments were performed in duplicate.

### Cytotoxicity of disquarimide compounds on BMDM

BMDM were infected with DsRed *L. major* parasites and cultured in the presence or absence of disquarimide compounds for 72 h at which time supernatants were collected. For a positive control to induce BMDM cell death, BMDMs were cultured with 10 μg/mL cycloheximide (CHX, Sigma) and 10 ng/mL recombinant mouse TNFα (Peprotech). To measure LDH released from non-viable cells using the LDH-Glo Cytotoxicity Assay (Promega), a small amount of culture medium (2–5 µL) was removed and diluted into LDH Storage Buffer at a dilution factor of 1:100 although the dilution factor can vary depending on the amount of cells and the presence of serum in the medium (Promega). LDH activity was measured by adding an equal volume of LDH Detection Reagent (50 µL) to the diluted sample (50 µL). Samples were incubated at RT for 60 minutes at which time luminescence was recorded using a CLARIOstar plate reader (BMG Labtech).

### Statistics

All data were analyzed using GraphPad Prism 9. Statistical significance was calculated using an unpaired 2-tailed Student’s for a single comparison between groups and *p* ≤ 0.05 was considered statistically significant. A Grubbs’ test was used to identify and mathematically remove outlier data points.

## RESULTS

Previous reports on both monosquaramide and disquaramide syntheses used a variety of solvents (diethyl ether, acetonitrile, or ethanol) and temperatures above room temperature ^12, 13, 17^. A major concern in generating safe and effective anti-*Leishmania* compounds is to greatly reduce the cost of the synthesis of the drug. To address this need, all commercial starting reagents for compound synthesis (3,4-diethoxycyclobut-3-ene-1,2-dione and the amines) in this study were low-cost and readily available through commercial means. The only solvent used for the syntheses was 200-proof ethanol, eliminating the need for peroxide-forming ethers and expensive acetonitrile. The compounds were all synthesized in a short two-step synthesis and all reactions were performed at ambient temperature without the use of catalysts or other additives to afford the target compounds in moderate to high yields. In addition, flash chromatography was only necessary for purification in a few cases (**3**, **24**, and **26**), the majority of the target compounds were isolated using simple vacuum filtration, greatly reducing the need for solvents.

Monosquaramides (**2** and **3**) were synthesized via an addition-elimination reaction of 3,4-diethoxycyclobut-3-ene-1,2-dione and the requisite amines in 200-proof ethanol in high yields (>80%) (Fig. 1). To control for the addition of only one amino side chain, strict stoichiometric controls were used (1:1 mole ratio of amine:3,4-diethoxycyclobut-3-ene-1,2-dione), the amines and 3,4-diethoxycyclobut-3-ene-1,2-dione were both diluted in ethanol, and the amines were added in slow, dropwise process to the 3,4-diethoxycyclobut-3-ene-1,2-dione solution. If any of these precautions were not undertaken, a significant amount of symmetrical disquaramide was synthesized. Once the monosquaramides were successfully synthesized, disquaramides were facily synthesized via another addition-elimination reaction using monosquaramides **2** or **3** with the requisite amines in 200-proof ethanol (Fig. 1). Reaction times were significantly reduced and percent yields were dramatically increased by adjusting the stoichiometry (2:1 mole ratio of amine:monosquaramide) of the reaction. Despite no further risk of accidental addition of amino groups, precautions with these reactions still needed to be maintained. Due to the exothermic nature of these reactions, amines were added in a dilute solution of ethanol in a slow, dropwise manner to a dilute solution of monosquaramides **2** or **3**. Using compound **17** as the lead compound ^12^, analogs **4**−**27** (Fig. 2) were successfully synthesized in moderate to high yields (48%−99%) in the hopes of finding a more potent antileishmanial compound as well as elucidating a more comprehensive structure-activity relationship.

**Figure 1.**
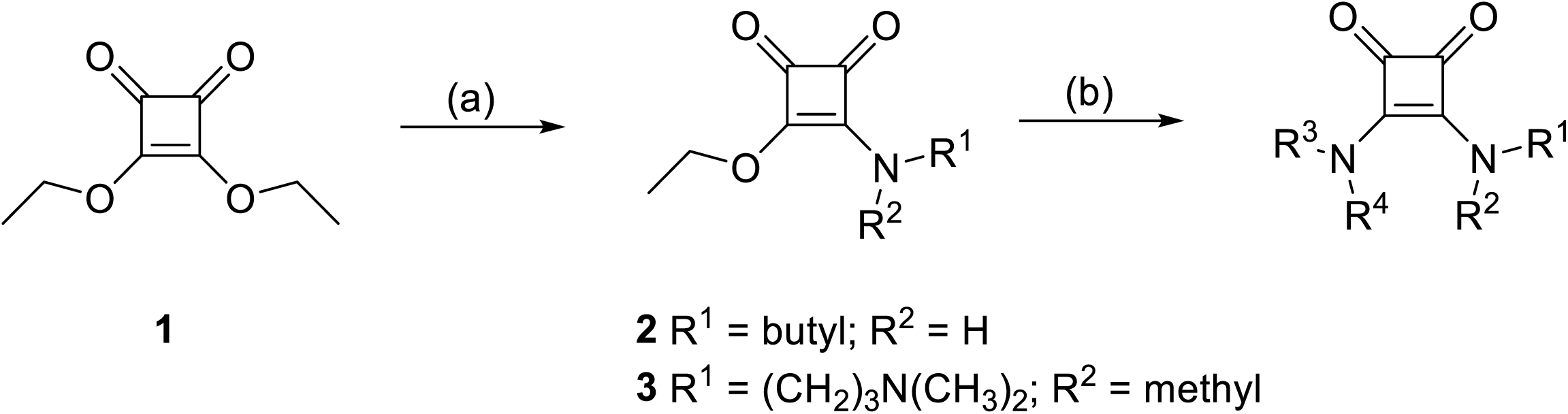
Scheme of synthesis of *N*,*N*-disquaramides. (a) *n*-butylamine (1.0 eq.) or *N*^1^,- *N*^1^,*N*^3^-trimethylpropane-1,3-diamine (1.0 eq.), EtOH (15 mL), room temperature, 2 h; (b) HNR_3_R_4_ (2.0 eq.), EtOH (15 mL), room temperature, 2−16 h).

**Figure 2.**
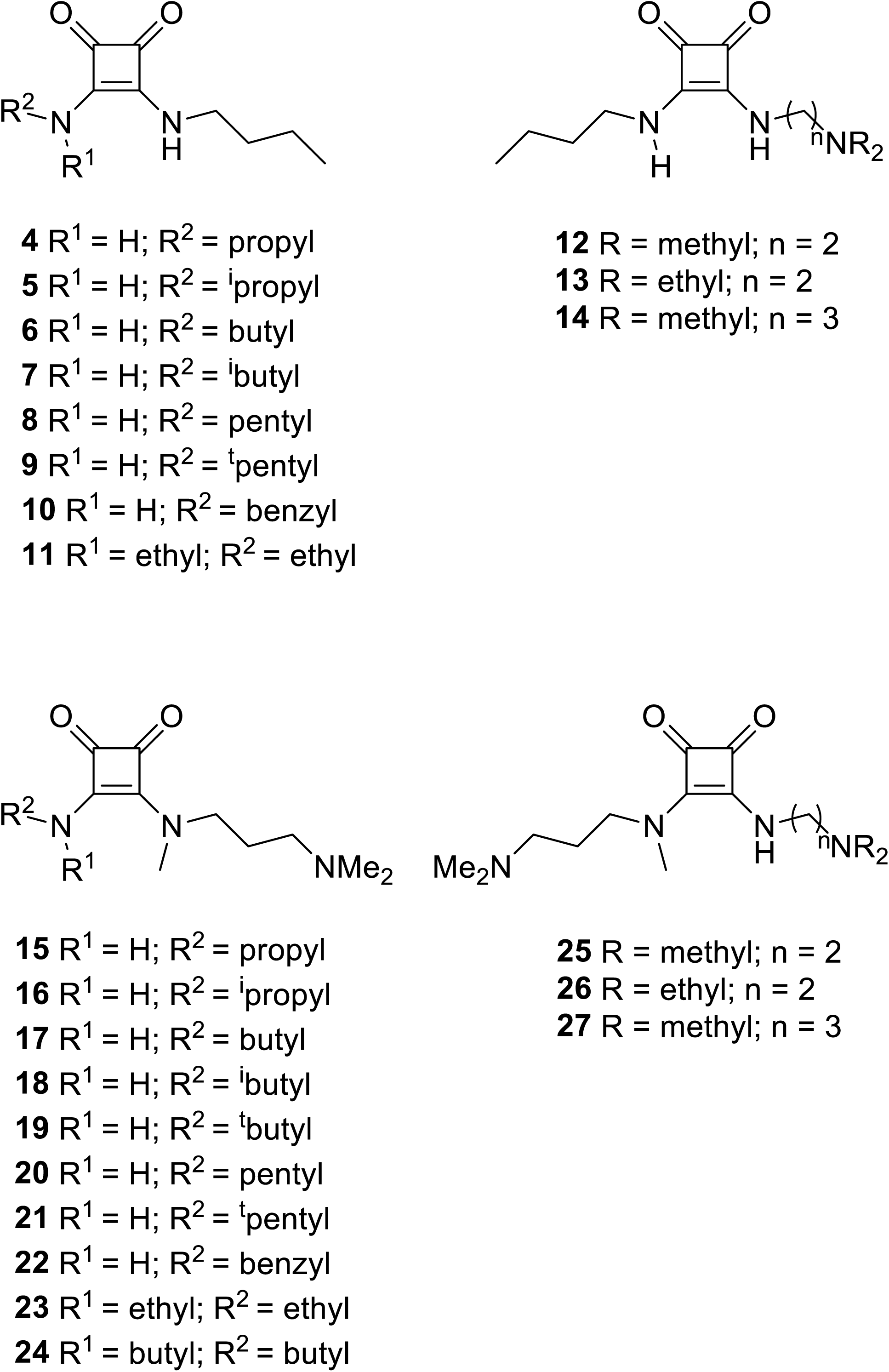
*N*,*N*’-Disquaramides evaluated against *L. major*.

The leishmanicidal activity of disquaramide compounds was tested on the promastigote stage of the parasite using fluorescently-labeled DsRed *L. major*. Using an in vitro assay, we show parasites increase in numbers during the first 4 days of culture (Fig. 3A and C). Acriflavine was used as a positive control for parasite killing as it eliminates kDNA through DNA intercalation ^18^. As expected, 1.0 and 2.0 mg/mL of acriflavine decreased parasites at days 3 and 4 compared to parasites cultured in media alone (Fig. 3B-C). To determine the effects of disquaramide compounds on *Leishmania* parasites, DsRed fluorescent parasites were cultured with 1, 5, or 10 μM of individual disquaramide compounds. Images of parasites cultured with 5 μM of the specified disqauramide compound at day 4 are shown (Fig. 3D and F). Quantification of parasite growth from days 1-4 cultured with or without the disquaramide compounds are reported (Fig. 3E and G). Upon incubation, compounds **13**, **20**, and **26** significantly decreased parasites after 4 days of culture at all three concentrations compared to control parasites which received no drug treatment (Fig. 3D-G). Compound **22** also inhibited parasite growth at both 5 and 10 μM concentrations of the compound (Fig. 3F-G). In contrast, disquaramide compounds **17**, **19**, and **24** were unable to slow parasite growth compared to parasites cultured in media alone without drug treatment (Fig. 3D-G). These data suggest that some disquaramide compounds can either perturb the growth of parasites or exhibit leishmanicidal effects, but not all disquaramide compounds are active against *L. major* parasites.

**Figure 3.**
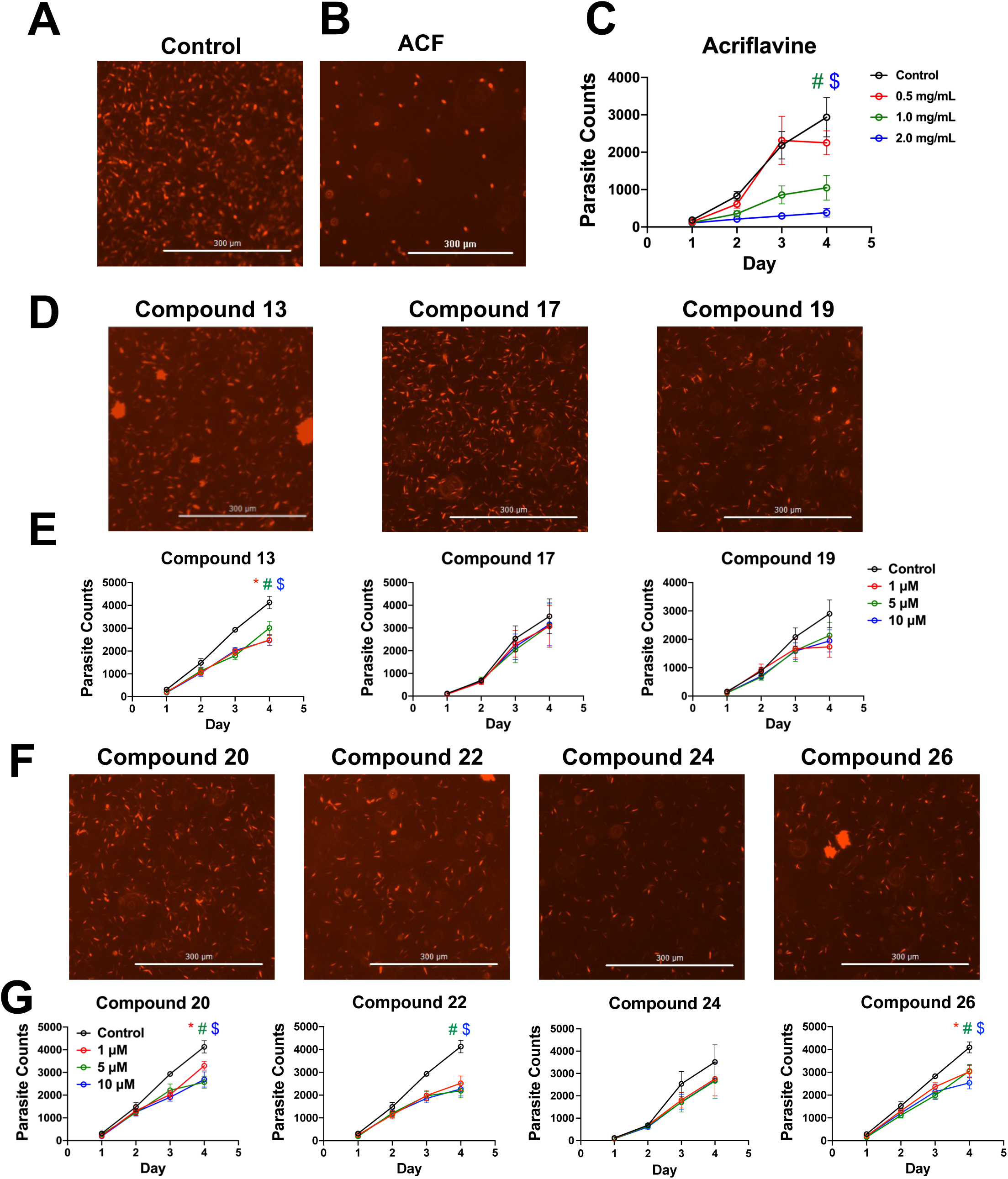
Disquaramide compounds compromise parasite viability. Parasites were cultured alone or in media containing 1, 5, or 10 μM of the specified disquaramide compound. At each time point, parasite viability was analyzed using a BioTek Cytation C10 Confocal Imaging Reader. **(A)** Images of parasites in media alone after 72 hours. **(B)** Images of parasites cultured with 2 mg/mL acriflavine after 72 hours. **(C)** The curve displays parasite growth without drug treatment and with treatment of acriflavine at 0.5, 1, and 2 mg/mL. **(D)** Parasite viability is shown in images taken of parasites cultured in 5 μM of the stated disquaramide compound. **(E)** Growth curves of parasites cultured in each disquaramide compound at a concentration of 1, 5, or 10 μM from day 1 to day 4. **(F)** Parasite viability is shown in images taken of parasites cultured in 5 μM of the stated disquaramide compound. **(G)** Growth curves of parasites cultured in each disquaramide compound at a concentration of 1, 5, or 10 μM from day 1 to day 4. Data are shown as mean ± SEM and significance (*p* ≤ 0.05) was determined using a t-test comparing each concentration against the control (red * denotes the significance of 1 μM compared to the control, green # denotes the significance of 5 μM against the control, and a blue $ denotes the significance of 10 μM compared to the control). Data are pooled from data of 3 independent experiments performed in triplicate. For images scale bar = 300 μm.

The previous experiments were conducted on parasites in culture alone without macrophages, but because parasites reside inside macrophages, we characterized the disquaramide-induced toxicity on the host, and the effects of disquaramide compounds on parasites residing in host cells. To test the effect of disquaramide compounds on cellular viability, BMDM were infected or not with DsRed *L. major* parasites. After parasite infection for 2 hours, extracellular parasites were washed away and cells were incubated with disquaramide compounds for 72 hours. An LDH assay on supernatants analyzed cell viability. Infected BMDM cultured in media alone was used as a negative control for LDH release. Cycloheximide in combincation with TNFα was used as a positive control to induce cell death for LDH release (Fig. 4A and B). Importantly, none of the disquaramide compounds elicited LDH release suggesting that these compounds are not cytotoxic for macrophages (Fig. 4A and B). Additionally, infection status did not affect LDH release because both uninfected and infected macrophages displayed similar LDH release compared to control macrophages (Fig. 4A and B).

**Figure 4.**
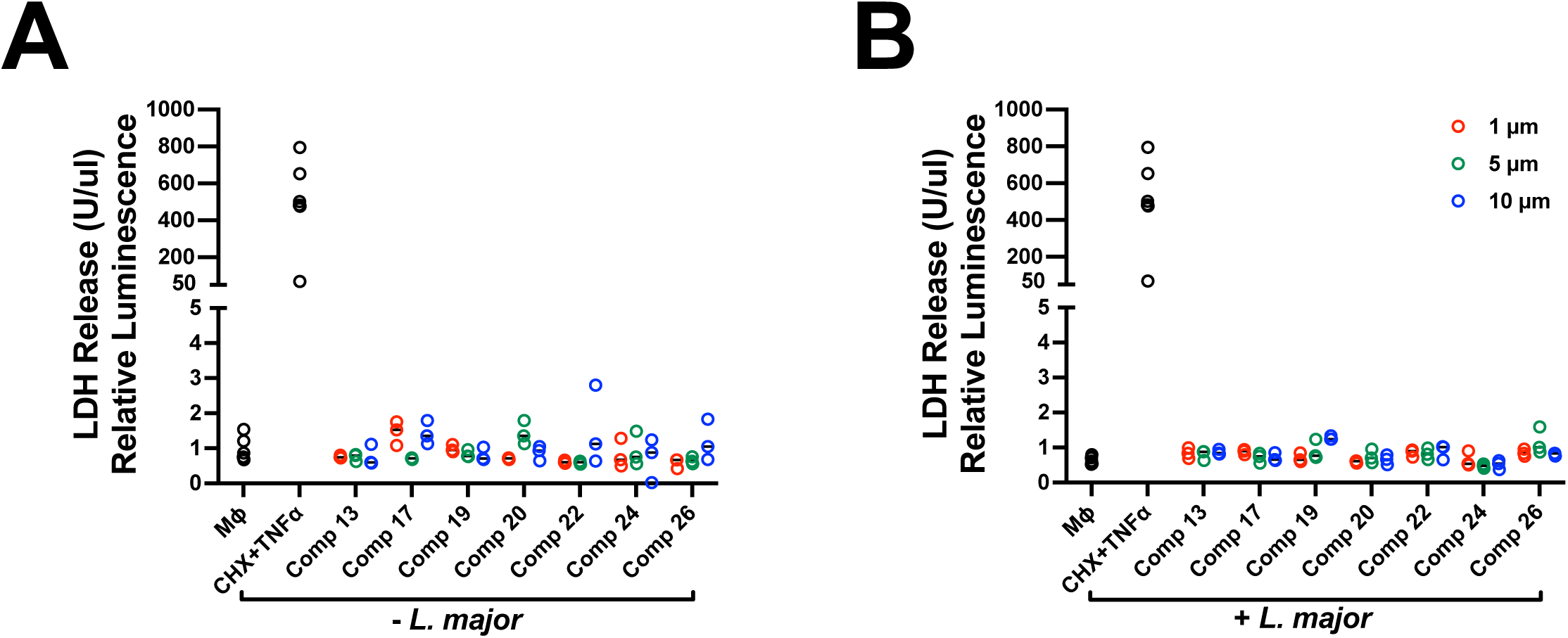
Disquaramide compounds do not induce host cell cytotoxicity. BMDM were infected with DsRed *L. major* parasites and cultured with or without disquaramide compounds for 72 hours at which point supernatants were collected. An LDH assay was conducted to measure cytotoxicity. **(A)** LDH release is shown in uninfected macrophages cultured with each disquaramide at 1, 5, or 10 μM. **(B)** LDH release from macrophages infected with DsRed *L. major* and cultured with 1, 5, or 10 μM of the disquaramide compounds. 10 μg/mL cycloheximide (CHX) and 10 ng/mL TNFα were used as a positive control for LDH release. Data are one representative experiment of at least 2 independent experiments performed with at least three wells per condition.

Next, the effect of disquaramide compounds on intracellular amastigotes was tested. As described above, BMDM were infected with DsRed *L. major* parasites and then cultured with or without disquaramide compounds for 72 hours. After 72 hours, infected macrophages were fixed and stained with DAPI to visualize the host cell. Figure 5 shows representative images of infected macrophages treated or not with various disquaramide compounds. For a negative control, infected BMDM were incubated without any compounds (media alone). First, we validated the semi-high-throughput assay to assess parasite burdens in macrophages. As expected, infected BMDM displayed significantly more parasites/macrophage than uninfected macrophages (Fig. 5A and C). Miltefosine is commonly used in the clinic to treat human leishmaniasis and was used as a positive control for parasite killing. Validating the assay, miltefosine decreased parasites at day 3 compared to parasites cultured in BMDMs in media alone (Fig. 5B and D). Miltefosine was effective at all concentrations tested including 1, 5, and 10 μM (Fig. 5D). Each disquaramide compound was capable of significantly decreasing the number of parasites/macrophage in at least one of the concentrations tested compared to control (Fig. 5E-H). Impressively, compound **24** significantly decreased the number of parasites/macrophage in all three concentrations tested compared to control infected macrophages (Fig. 5G-H). Compounds **20** and **22** significantly decreased parasite numbers at concentrations of 1 μM and 5 μM, but not 10 μM (Fig. 5G-H). Compound **17** inhibited parasites at concentrations of 5 μM and 10 μM but not 1 μM (Fig. 5E-F). Compounds **13**, **19**, and **26** only decreased parasite counts at one concentration: **13** (5 μM), **19** (5 μM), and **26** (10 μM) (Fig. 5E-H). Taken together, these results show disquaramide compounds are effective anti-leishmanial agents against intracellular parasites residing in macrophages.

**Figure 5.**
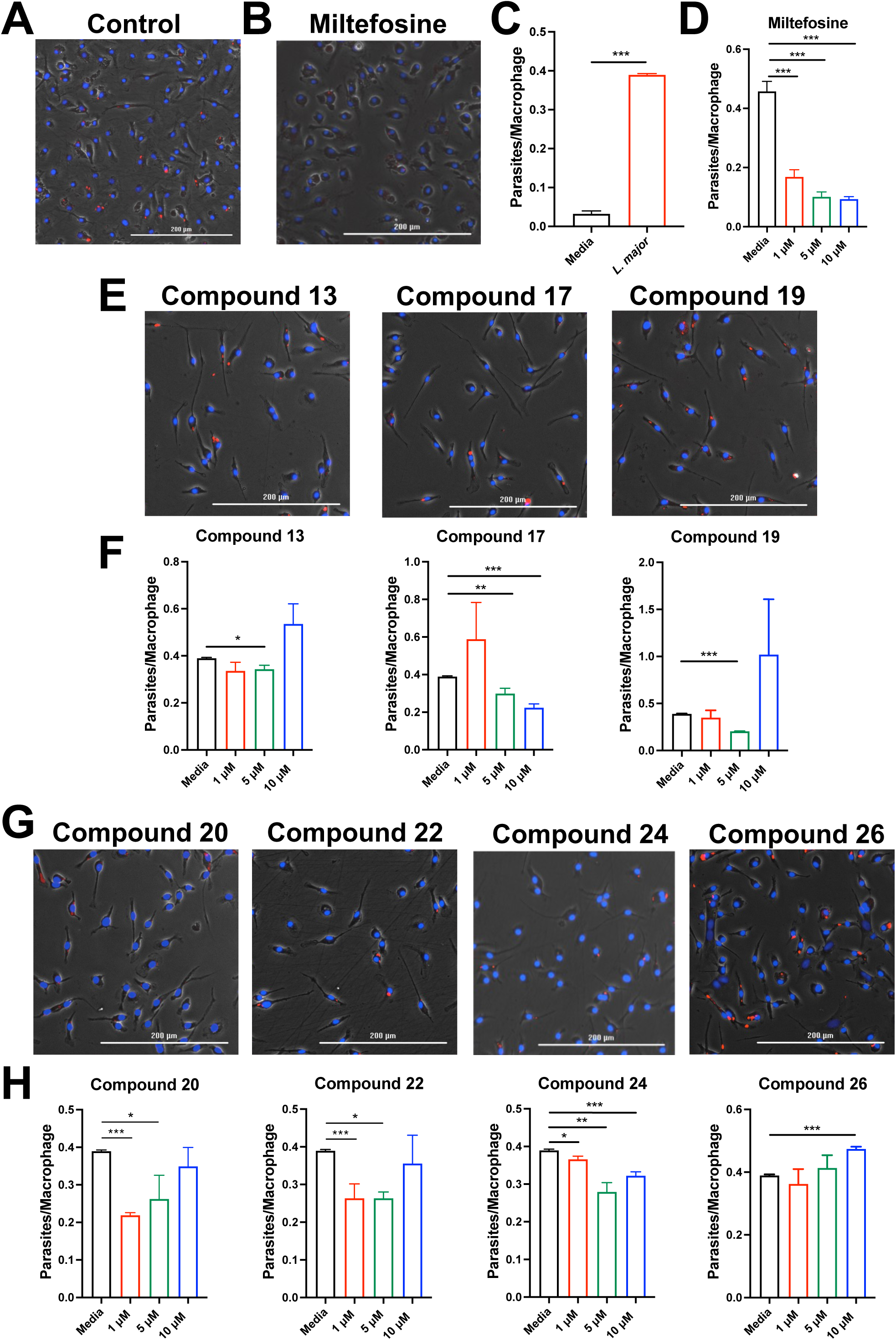
Intracellular *L. major* parasites are sensitive to disquaramide compound treatment. BMDM were infected or not with DsRed *L. major* parasites at an MOI of 5:1. Infected BMDM were cultured in the presence of various disquarimide compounds for 72 hours. Cells were fixed and stained with DAPI. Intracellular parasites were imaged and quantified using BioTek Cytation C10 Confocal Imaging Reader. **(A)** Images of DsRed *L. major* parasites in BMDM cultured in media alone after 72 hours. **(B)** Images of parasites in BMDM cultured in 5 μM miltefosine after 72 hours. Miltefosine is used as a positive control for parasite killing. **(C)** Quantification of DsRed *L. major* in images in A showing the number of parasites per macrophage. Uninfected BMDM (media), infected untreated BMDM, and infected BMDM treated with 10 μg/mL CHX and 10 ng/mL TNFα show the specificity of the assay to quantify the number of parasites per macrophage. **(D)** Quantification of DsRed *L. major* in images in B showing the number of parasites per macrophage after 72 hours of 1, 5, or 10 μM miltefosine. **(E)** Images of DsRed *L. major* parasites in BMDM cultured in 5 μM of stated disquaramide compound after 72 hours. **(F)** Each disquaramide compound was tested at concentrations of 1, 5, and 10 μM. **(G)** Images of DsRed *L. major* parasites in BMDM cultured in 5 μM of stated disquaramide compound after 72 hours. **(H)** Each disquaramide compound was tested at concentrations of 1, 5, and 10 μM. Data are shown as mean ± SEM. Statistical significance (*p* ≤ 0.05) between parasites/macrophage after varying concentrations of drug treatment compared to controls determined using a t-test. \**p* ≤ 0.05, \*\**p* < 0.01, \*\*\**p* < 0.001, *t-*test comparing uninfected versus infected or comparing control media to drug treatment. Data are one representative experiment of 3 independent experiments performed in triplicate. For images scale bar = 200 μm.

## DISCUSSION

In vitro testing of disquaramide compounds reveals that they exhibit anti-leishmanial activity against both parasites themselves as well as parasites inside infected host cells. Importantly, disquaramide compounds exhibit little to no toxicity against host cells. The short synthesis time, low-cost of starting reagents, and simple purification methods have led to disquaramide compounds being investigated as low-cost drugs against parasites causing NTDs including leishmaniasis and Chagas disease, as well as multiple bacterial strains including MRSA ^19^. Previous studies on disquaramide compounds against *Leishmania* indicate that the primary route of parasite killing is by damage to the parasite membranes leading to widespread damage to overall structure as well as to individual organelles, often rendering the parasites unrecognizable ^12^. Other studies suggest that disquaramide compounds interfere with the function of ATP synthase on *Mycobacterium* ^20^. While we did not investigate the structural integrity of parasite membranes by electron microscopy or measure ATP synthase function, the similar structure, and effectiveness of our tested disquaramides to those tested in previous studies suggest a similar method of killing ^12^.

Despite their effectiveness against *Leishmania* parasites, no toxic effects were observed in BMDM treated with disquaramides at any concentrations tested. The observations that disquaramide compounds reduce parasite numbers, directly in parasite cultures and when parasites reside in host macrophages, also suggest that the antileishmanial effect of the disquaramide compounds is independent of the host. Our findings are consistent with previous reports using disquaramide compounds against *Leishmania* which also showed little to no toxicity against host cells ^12^. The lack of host cell cytotoxicity bodes well for testing their effectiveness in future in vivo studies and possible therapeutic applications.

Generally, our results found that compounds containing smaller hydrophobic substituent arms (**15, 16, and 18**) as well as two hydrophobic substituent arms (**4−11**) proved ineffective against *L. major* parasites. Looking at the structural motifs of the more successful compounds that were tested, **13**, **19**, **20**, **22**, and **24**, each contain a bulkier hydrophobic arm (butyl, *t*-butyl, pentyl, benzyl, and di-*n*-butyl, respectively) and a more hydrophilic diamino arm (*N*^1^,*N*^1^,*N*^2^-trimethylpropaminediamine (**19**, **20**, **22** and **24**) or *N*^1^,*N*^1^-diethylethylenediamine (**13**). The lone exception to this trend is **26,** which contains two hydrophilic diamino arms (*N*^1^,*N*^1^,*N*^2^-trimethylpropaminediamine and *N*^1^,*N*^1^-diethylethylenediamine), making it unique compared to the other successful compounds.

A combination of a hydrophobic butyl arm and a more hydrophilic diamino *N*^1^,*N*^1^,*N*^2^-trimethylpropaminediamine arm, **17**, was our lead compound ^12^. Our results reinforce the benefits of designing disquaramide compounds that contain both a larger, hydrophobic arm (alkyl or aryl) and a more hydrophilic diamino arm. Additionally, the realization of the larger, and more hydrophobic arms and alternative hydrophilic diamino arms as viable, and more potent, substituents opens up a much more targeted second generation of compounds. Of particular note, installing aryl groups, as seen in compound **22**, as well as *N*^1^,*N*^1^-diethylethylenediamine, as seen in compounds **13** and **26**, opens up two relatively unexplored disquaramide substituents. These future gerenerations of compounds will explore an increase in hydrophobicity in one arm as well as more variety of the diamino substituent arm.

## CONCLUSION

A drug library consisting of 24 disquaramide compounds was successfully synthesized and tested in vitro against *L. major* parasites directly and inside host macrophages. In general, the synthesized disquaramides demonstrated effectiveness against *Leishmania* parasites in both cases, indicating that disquaramides have potential as leishmanicidal agents. Additionally, these compounds showed little to no toxicity to the host cells. Of the 24 disquaramides tested, six of them demonstrated the most potential: **13**, **19**, **20**, **22**, **24**, and **26**. Compounds **13** and **22** are of particular interest due to their relatively unexplored substituents in regard to anti-*Leishmania* activity, thus opening up an avenue for further structural optimization in the creation of a next-generation disquaramide drug library.

### Limitations

Most of the disquaramide compounds did not exhibit a dose-dependent effect on parasite numbers either by inhibiting parasite growth or inducing death. We only used BMDM as a part of this screen of disquaramide compounds and in the future we will use other types of primary macrophages, such as peritoneal macrophages to test the leishmanicidal effects of disquaramide compounds. This initial screen of disquaramide compounds on *Leishmania* parasites was limited to *L. major*, but future studies will examine the effects of other parasite species, such as *L. mexicana*, and *L. amazonensis*, and *L. braziliensis*. Lastly, this study focused on characterizing the effects of disquaramide compounds on parasites directly as well as parasites residing inside macrophages in vitro, but additional in vivo experiments need to be performed testing the in vivo activity of disquaramide compounds using our preclinical *Leishmania* mouse model of infection (parasite inoculation in the ear of C57BL/6 mice) ^21^.

## SUPPORTING INFORMATION

The supporting information contains the ^1^H and ^13^C NMR spectra of all previously unreported compounds (PDF).

## ACKNOWLEDGMENTS

This work was supported by the Arkansas IDeA Network for Biomedical Research Excellence (INBRE) (funded by the National Institutes of Health (NIH) National Institute of General Medical Sciences (NIGMS) Centers of Biomedical Research Excellence Grant P20-GM103429) and the Center for Microbial Pathogenesis and Host Inflammatory Responses (funded by NIH NIGMS Centers of Biomedical Research Excellence Grant P20-GM103625). This publication was also supported in part by funds provided by the National Center For Advancing Translational Sciences of the NIH under awards TL1 TR003109 and UL1 TR003107 for the Systems Pharmacology and Therapeutics (SPaT) NIH T32 training grant GM106999 to Lucy Fry. This work was also supported by the Arkansas INBRE PRO Program summer internship (funded by NIH NIGMS Grant P20-GM103429) to Alexx Weaver. On a more personal note, the authors would like to thank Dr. Larry Cornett, Dr. Jerry Ware, and Caroline Miller for their endless efforts to strengthen academic research throughout Arkansas and for introducing Drs. Weinkopff and Naumiec. The content is solely the responsibility of the authors and does not necessarily represent the official views of the NIH. The funders had no role in study design, data analysis, decision to publish, or preparation of the manuscript.

